# Detecting tropical selective logging with SAR data requires a time series approach

**DOI:** 10.1101/2020.03.31.018606

**Authors:** MG Hethcoat, JMB Carreiras, DP Edwards, RG Bryant, S Quegan

## Abstract

Selective logging is the primary driver of forest degradation in the tropics and reduces the capacity of forests to harbour biodiversity, maintain key ecosystem processes, sequester carbon, and support human livelihoods. While the preceding decade has seen a tremendous improvement in the ability to monitor forest disturbances from space, advances in forest monitoring have almost universally relied on optical satellite data from the Landsat program, whose effectiveness is limited in tropical regions with frequent cloud cover. Synthetic aperture radar (SAR) data can penetrate clouds and have been utilized in forest mapping applications since the early 1990s, but no study has exclusively used SAR data to map tropical selective logging. A detailed selective logging dataset from three lowland tropical forest regions in the Brazilian Amazon was used to assess the effectiveness of SAR data from Sentinel-1, RADARSAT-2 and PALSAR-2 for monitoring tropical selective logging. We built Random Forest models in an effort to classify pixel-based differences in logged and unlogged areas. In addition, we used the BFAST algorithm to assess if a dense time series of Sentinel-1 imagery displayed recognizable shifts in pixel values after selective logging. Random Forest classification with SAR data (Sentinel-1, RADARSAT-2, and ALOS-2 PALSAR-2) performed poorly, having high commission and omission errors for logged observations. This suggests little to no difference in pixel-based metrics between logged and unlogged areas for these sensors. In contrast, the Sentinel-1 time series analyses indicated that areas under higher intensity selective logging (> 20 m^3^ ha^−1^) show a distinct spike in the number of pixels that included a breakpoint during the logging season. BFAST detected breakpoints in 50% of logged pixels and exhibited a false alarm rate of approximately 10% in unlogged forest. Overall our results suggest that SAR data can be used in time series analyses to detect tropical selective logging at high intensity logging locations within the Amazon (> 20 m^3^ ha^−1^). These results have important implications for current and future abilities to detect selective logging with freely available SAR data from SAOCOM 1A, the planned continuation missions of Sentinel-1 (C and D), ALOS PALSAR-1 archives (expected to be opened for free access in 2020), and the upcoming launch of NISAR.

## 1. Introduction

Selective logging is the primary driver of forest degradation in the tropics (Curtis et al., 2018; Hosonuma et al., 2012). Logging reduces the capacity of forests to harbour biodiversity, maintain key ecosystem processes, sequester carbon, and support human livelihoods (Baccini et al., 2017; Barlow et al., 2016; Lewis et al., 2015). However, large uncertainties remain in assessing the true impact of selective logging because the technological advances in detecting and monitoring logging at large scales are only just emerging (Hethcoat et al., 2019). The ability to reliably map forest degradation from selective logging is a key element in understanding the terrestrial portion of the carbon budget and the role of land-use in turning tropical forests into net carbon emitters (Baccini et al., 2017). In addition, reliable forest monitoring systems are urgently needed for tropical nations and conservation groups seeking to report and/or mitigate carbon emissions through improved forest stewardship (GOFC-GOLD, 2016).

While the preceding decade has seen a tremendous improvement in the ability to detect forest disturbances from space (Hansen et al., 2013; Hethcoat et al., 2019; Tyukavina et al., 2017), advances in forest monitoring have almost universally relied on optical satellite data from the Landsat program. Yet, the effectiveness of optical data is limited in tropical regions with frequent cloud cover like the northwest Amazon and central Africa. Synthetic aperture radar (SAR) data can penetrate clouds and have been utilized in forest mapping applications since the early 1990s (reviewed in Koch, 2010). However, the SAR data archives are spatially and temporally fragmented, and in many cases the data products required commercial licences for their use. Consequently, uptake by users has been more limited than optical data and the full potential of SAR has likely been under-utilized (Reiche et al., 2016).

SAR backscatter, particularly at L- and P-band, is sensitive to changes in carbon stocks in forests with biomass < 300 Mg ha^−1^ (Koch, 2010; Mitchard et al., 2009; Saatchi et al., 2011). This enables accurate differentiation between forested and non-forested areas and has been well studied (e.g. Shimada et al., 2014). More recently, polarimetric and interferometric methods have been developed that utilize information in the SAR signal to detect forest changes (Deutscher et al., 2013; Flores-Anderson et al., 2019; Lei et al., 2018; Mathieu et al., 2013). Yet, the limited temporal and spatial coverage of SAR data have hampered widespread application and use of these techniques to monitor forest disturbances (e.g. single-pass interferometric SAR is only available with TanDEM-X data). Moreover, advancements in monitoring selective logging with SAR data are generally lacking, despite widespread recognition of both the need and the role it could play (Mitchell et al., 2017; Reiche et al., 2016).

The launch of Sentinel-1A in mid-2014 represented the first continuous global acquisition strategy for open SAR data. Since that time two studies have exclusively used Sentinel-1 to map deforestation (Antropov et al., 2016; Delgado-Aguilar et al., 2017), with others utilizing a fusion of optical and SAR data (Joshi et al., 2016; Reiche et al., 2018a, 2018b, 2015). While methods that fuse optical and Sentinel-1 have been successful, their continued dependence on optical imagery nevertheless limits their utility in regions with frequent cloud cover. With the successful launch of SAOCOM 1A in late 2018, the planned continuation of the Sentinel-1 missions (with C and D), and the anticipated launches of SAOCOM 1B in 2019 and NISAR in 2021, vast amounts of free C- and L-band SAR data will soon be available. Accordingly, methods are needed that utilize SAR data for large-scale forest monitoring, yet no study has used Sentinel-1 for detection of selective logging activities.

The primary objective of this paper was to assess the ability of Sentinel-1 to detect tropical selective logging. Detailed spatial and temporal logging records from three regions in Brazil were used to develop and test the effectiveness of two different detection techniques: (1) exploiting pixel-based differences between logged and unlogged locations in single images and (2) detecting change in a time series of pixels known to be logged.

Pixel-based methods for detecting changes in remotely sensed imagery often utilize differences between pixel values or other mathematically derived metrics in time or space, for example before and after some disturbance or in areas known to be disturbed and undisturbed within the same image (reviewed in Hussain et al., 2013). These differences can be used for classification, employed in machine learning, or analyzed temporally to map change. Recently, the detection of selectively logged regions in single images has been demonstrated successfully with optical data from Landsat (Hethcoat et al. 2019). Accordingly, we sought to evaluate whether similar methods could be transferred to SAR data. The selective logging records were used to build supervised machine learning models to detect selective logging. Machine learning methods have many applications in remote sensing and have been used with increasing frequency and success (Lary et al., 2018). We performed equivalent analyses with SAR data from the C-band RADARSAT-2 and L-band PALSAR-2 sensors to compare the performance of longer wavelength (i.e. L-band PALSAR-2) and higher resolution data (both RADARSAT-2 and PALSAR-2 have higher sensor resolution).

In addition, we used all the available Sentinel-1 archives in a time series analysis to monitor pixel values for breakpoints in the time series of locations that had been selectively logged. Time series methods have increasingly been used for monitoring changes in pixel values, in part because of the availability of vast archives of imagery on cloud computing platforms like Google Earth Engine (Gorelick et al., 2017), but also because of the recognition that seasonal or longer term trends in pixel values can be less susceptible to erroneously characterizing change (Bullock et al., 2018; Verbesselt et al., 2012; Zhu, 2017). Given that forest disturbances from selective logging are often subtle and short-lived, detecting changes with SAR data over large regions will present technological and algorithmic challenges. However, a critical assessment of detection capabilities and a careful understanding of the performance of these data types is essential for advancing forest monitoring techniques in the tropics.

## 2. Study area and data

### 2.1. Study area and selective logging data

Selective logging data from three lowland tropical forest regions in the Brazilian Amazon were used in this study (Figure 1). The Jacunda and Jamari regions are inside the Jacundá and Jamari National Forests, Rondônia, while the Saraca region is inside the Saracá-Taquera National Forest, Pará. Forest inventory data from 14 forest management units (FMUs) selectively logged between 2012 and 2017 were used, comprising over 32,000 individual tree locations. Unlogged data from three additional locations, one inside each study region, comprised over 11,500 randomly selected point locations known to have remained unlogged during the study period (Table 1).

**Table 1.** Data used in the classification of selective logging from three study regions in the Brazilian Amazon. The forest management unit (FMU), logging intensity, sample size (pixels), and overlap with satellite data coverage are shown for Sentinel-1 (S), RADARSAT-2 (R), and PALSAR-2 (P).

| FMU | Logging Intensity<br>(m<sup>3</sup> ha<sup>-1</sup>) | N | Coverage |
| --- | --- | --- | --- |
| Jacunda_I_2016 | 6 | 2,290 | S |
| Jacunda_I_2017 | 9 | 2,822 | S |
| Jacunda_II_2015 | 15 | 2,613 | S |
| Jacunda_II_2016 | 10 | 1,815 | S |
| Jacunda_II_2017 | 7 | 1,310 | S, R<sup>*</sup> |
| Jacunda_Reserve | 0 | 3,000 | S<sup>*</sup>, R<sup>*</sup> |
| Jamari_I_2015 | 22 | 1,094 | S, R<sup>*</sup> |
| Jamari_I_2016 | 10 | 653 | S, R<sup>*</sup> |
| Jamari_I_2017 | 12 | 911 | S, R<sup>*</sup> |
| Jamari_III_2012 | 10 | 3,071 | R |
| Jamari_III_2015 | 11 | 3,042 | S, R<sup>*</sup> |
| Jamari_III_2016 | 9 | 2,058 | S, R<sup>*</sup>, P |
| Jamari_III_2017 | 11 | 2,597 | S, R<sup>*</sup>, P |
| Jamari_Reserve | 0 | 5,912 | S<sup>*</sup>, R<sup>*</sup>, P<sup>*</sup> |
| Saraca_Ia_2017 | 12 | 3,769 | S |
| Saraca_II_2016 | 25 | 3,223 | S |
| Saraca_II_2017 | 21 | 4,729 | S |
| Saraca_Reserve | 0 | 3,000 | S<sup>*</sup> |

<sup>\*</sup> FMU was unlogged at time of acquisition and data represent unlogged observations

**Figure 1.**
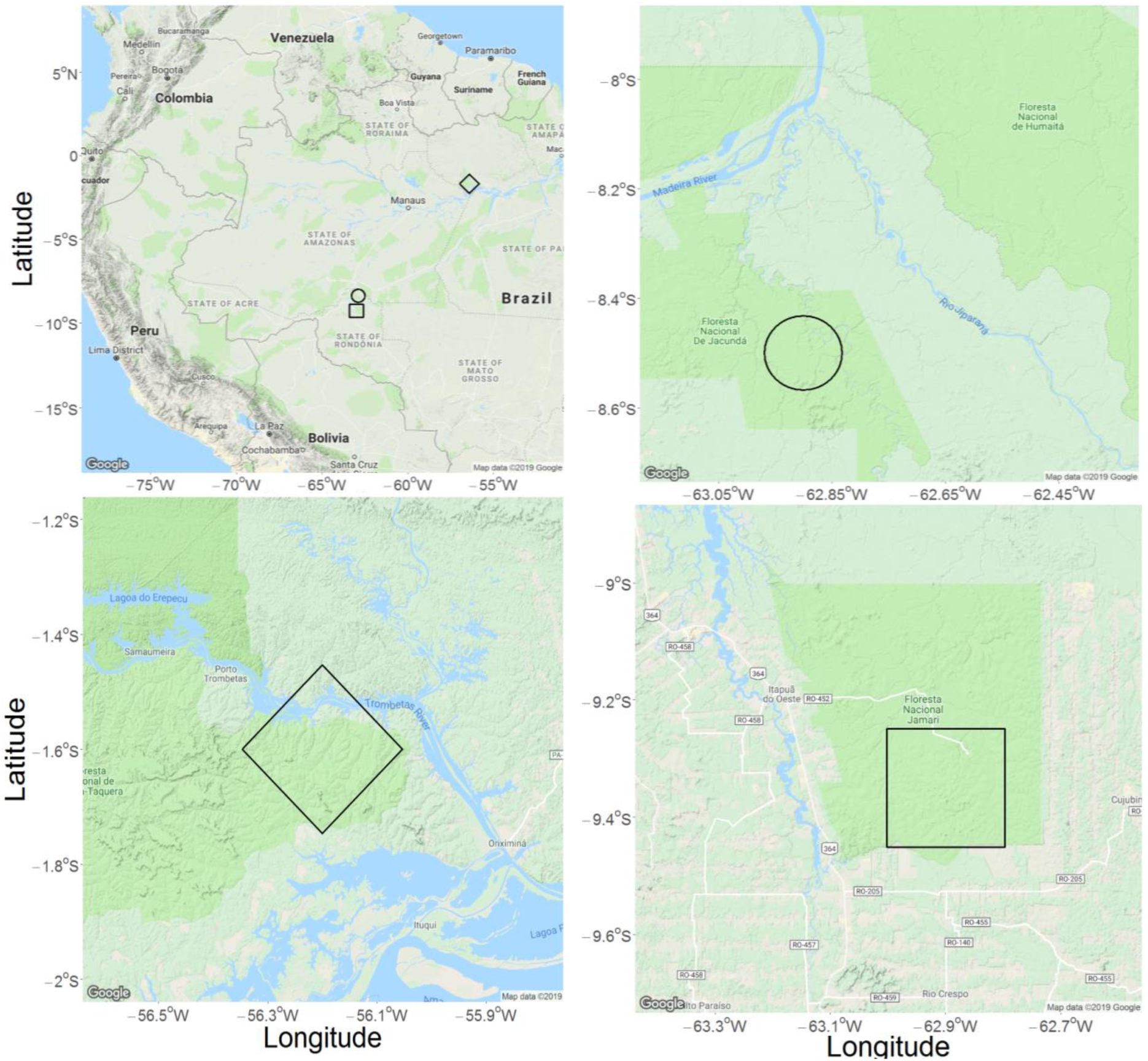
Location of the Jacunda (circle), Jamari (square), and Saraca (diamond) study regions in the Brazilian Amazon.

### 2.2. Satellite data and pre-processing

All available C-band Sentinel-1A Ground Range Detected scenes in descending orbit and Interferometric Wide mode (VV and VH) were utilized in Google Earth Engine (GEE) over the study regions through November 2018. These had incidence angles of 38.7°, and 38.7°, and 31.4° for Jacunda, Jamari and Saraca, respectively. GEE is a cloud computing platform hosting calibrated, ortho-corrected Sentinel-1 scenes that have been processed in the following steps using the Sentinel-1 Toolbox: (1) thermal noise removal; (2) radiometric calibration; and (3) terrain correction using the Shuttle Radar Topography Mission (SRTM) 30 m digital elevation model (DEM). The resulting images had a pixel size of 10 m.

Single Look Complex C-band RADARSAT-2 scenes in Fine mode (HH and HV) were obtained from the Canadian Space Agency. Twelve ascending scenes, with an incidence angle of 30.7°, coincided with selective logging records and were acquired between 2011 and 2012. Pre-processing of images was done with the Sentinel-1 Toolbox and included: (1) radiometric calibration; (2) multi-looking (by a factor of 2 in azimuth) to produce square pixels; and (3) terrain correction using the SRTM 30 m DEM. The resulting images had a pixel size of 10 m.

Level 2.1 L-band PALSAR-2 scenes (HH and HV) were obtained from the Japan Aerospace Exploration Agency (JAXA) with a pixel size of 6.25 m. Four geometrically corrected scenes coincided with selective logging records and were acquired between 2016 and 2017 with incidence angles of 28.5° in ascending orbit. Image digital number was converted to normalized backscatter using the calibration factors provided by JAXA.

### 2.3. Speckle filtering

SAR data are inherently speckled from interference between scattering objects on the ground (Woodhouse, 2017) and often require reduction of speckle prior to analyses. Many speckle-reduction methods involve spatial averaging, but the associated loss of spatial resolution was likely to hinder the detection of the subtle signal from selective logging activities. Thus, following the SAR pre-processing steps detailed above for each data type, the final step involved multi-temporal filtering to reduce speckle (Quegan and Yu, 2001). Multi-temporal filtering reduces speckle by averaging a pixel’s speckle through time (as opposed to a spatial average). A 7 ×7 pixel window was used. The equivalent number of looks after speckle filtering for Sentinel-1, RADARSAT-2 and PALSAR-2 was approximately 15, 5 and 5, respectively.

## 3. Methods

### 3.1. Supervised classification with Random Forest

#### 3.1.1. Data inputs for classifying selective logging

For each satellite data type (Sentinel-1, RADARSAT-2, and PALSAR-2) data were extracted at each pixel where logging occurred and randomly selected pixels in nearby regions that remained unlogged. Thus, the data inputs for logged and unlogged observations came from a single scene for each study region (i.e. a space-for-time study design in contrast to images before and after logging from the same location). Selective logging at the study areas only occurred during the dry season, approximately June-October in a given year, and data were extracted from images acquired as late into the logging period as possible (Table S1) to ensure the majority of pixels had been subjected to logging, but also before the onset of the rainy season (Hethcoat et al., 2019). In addition, logging activities tend to be accompanied by surrounding disturbances (canopy gaps, skid trails, patios, and logging roads) resulting in forest disturbances beyond just the pixels where a tree was removed. Accordingly seven texture measures were calculated for each polarization (sum average, sum variance, homogeneity, contrast, dissimilarity, entropy, and second moment) to provide a local context for each pixel (Haralick et al., 1973). These were calculated within a 7×7 pixel window, chosen as a trade-off between minimizing window size while still capturing the variability in selectively logged forests compared to unlogged forests. Finally, a composite band was calculated as the ratio of the co-polarized channel to the cross-polarized channel (i.e. HH/HV or VV/VH). Each dataset thus comprised a 17-element vector (2 polarization bands, their ratio composite band, and 7 texture measures for each polarization) for each pixel where logging occurred and randomly selected pixels that remained unlogged.

#### 3.1.2. Random Forests for classification of selective logging

We built Random Forest (RF) models using the *randomForest* package in program R version 3.5.1 (Liaw and Wiener, 2002; R Development Core Team, 2018). The RF algorithm (Breiman, 2001a) is an ensemble learning method for classification. Each dataset was split into 75% for training and 25% was withheld for validation. In order to further ensure the independence of training and validation datasets, the validation data were spatially filtered such that no observations in the training dataset were within 90 m of an observation in the validation dataset. RF models have two tuning parameters: the number of classification trees grown (*k*), and the number of predictor variables used to split a node into two sub-nodes (*m*). We used a cross-validation technique to identify the number of trees and the number of variables to use at each node that minimized the out-of-bag error rate on each training dataset (Table S2). The importance of each predictor variable was assessed during model training, using Mean Decrease in Accuracy, defined as the decrease in classification accuracy associated with not utilizing that particular input variable for classification (Breiman, 2001b).

#### 3.1.3. Model validation: assessing accuracy

RF models were validated using a random subset of the full dataset for each sensor (described in Section 3.1.2). By default, RF models assign an observation to the class indicated by the majority of decision trees (Breiman, 2001a). However, the proportion of trees that voted for a particular class from the total set of trees can be obtained for each observation and a classification threshold can be applied to this proportion (Hethcoat et al., 2019; Liaw and Wiener, 2002). We adopted such an approach, wherein the proportion of trees that predicted each observation to be logged, informally termed the *likelihood* a pixel was logged, was used to select the classification threshold. A threshold, *T*, was defined such that if *likelihood* > *T* the pixel was classified as logged (Figure 2).

**Figure 2.**
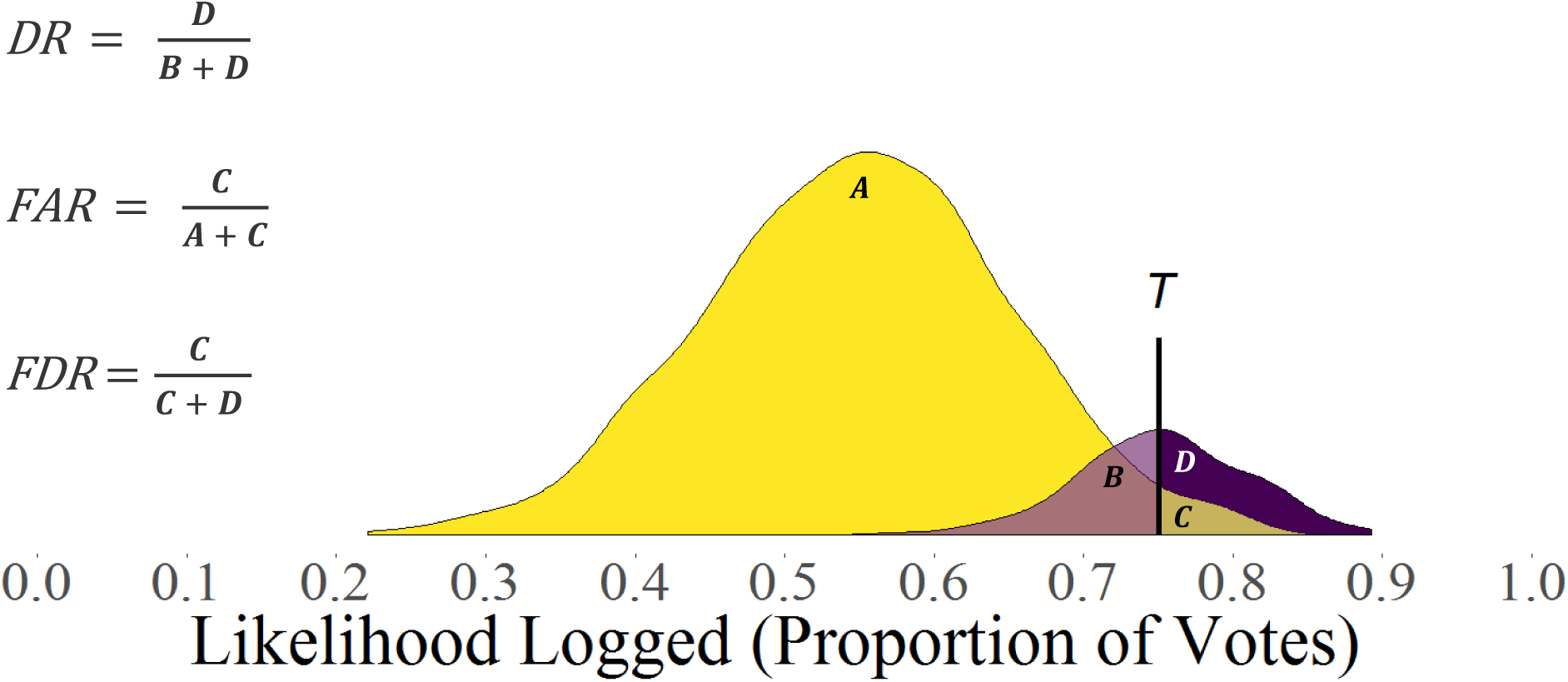
Diagram representing the trade-off between the detection rate (DR) and the false alarm rate (FAR) associated with using a threshold *T* (vertical black line) to label pixels as logged and unlogged based upon the proportion of votes that each observation was predicted to be logged. The purple and yellow colors correspond to density plots for hypothetical logged and unlogged observations, respectively. Thus, the areas A and B are the portions of the observations from unlogged and logged pixels, respectively, that will be labelled as unlogged. Similarly, C and D represent the portions of the observations from logged and unlogged pixels, respectively, that will be labelled as logged.

The confusion matrix then has the form:

|  |  | Reference |  |
| --- | --- | --- | --- |
|  |  | L | UL |
| Predicted | L | $D_L$ | $D_{UL}$ |
| | UL | $N_L - D_L$ | $N_{UL} - D_{UL}$ |

where L and UL refer to logged and unlogged classes, *N*_*L*_ and *N*_*UL*_ are the numbers of logged and unlogged observations in the reference dataset, and *D*_*L*_ and *D*_*UL*_ are the numbers of logged and unlogged pixels detected as logged, respectively. We defined the *detection rate DR = D*_*L*_*/N*_*L*_ and *false alarm rate FAR = D*_*UL*_*/N*_*UL*_ as the frequency that a logged or unlogged pixel was classified as logged, respectively. Thus, the DR is equivalent to 1 minus the omission error of the logged class and the FAR is the omission error of the unlogged class. In addition, we defined the *false discovery rate* (FDR):

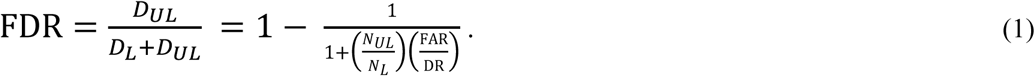

The FDR is the proportion of all observations that were detected as logged that were actually unlogged, and is equivalent to the commission error of the logged class. The FDR is an assessment of the rate of prediction error (i.e. type I) when labelling pixels as logged and can be used in detection problems with rare events or unbalanced datasets, such as selectively logged pixels within the Amazon Basin (Benjamini and Hochberg, 1995; Hethcoat et al., 2019; Neuvial and Roquain, 2012). A high DR and low FDR is clearly desirable, but these cannot be fixed independently in two-class detection problems and both depend on the threshold value (Figure 2). For example, if achieving a 95% detection rate led to a FDR of 50%, then half of all predictions of logging would be incorrect. This level of performance would make estimates of selective logging extremely uncertain. The value of the classification threshold (*T*) therefore represents a trade-off between true and false detections. In practice, a viable detection method would expect to achieve a DR > 50% while limiting the FDR to 10-20% to have any value for widespread forest monitoring. The performance of each sensor was assessed by plotting the DR, FAR and FDR values as *T* varied from 0 to 1 to facilitate discussion of model performance.

#### 3.1.4. Sentinel-1 classification of high intensity logging

Most of the selective logging data in this study were low-intensity (<15 m^3^ ha^−1^) and we anticipated the logging signal to be weak and difficult to detect. Consequently, we also considered a reduced Sentinel-1 dataset that included only those FMUs with logging intensities above 20 m^3^ ha^−1^ (*n* = 3 sites) and the unlogged data (*n* = 3 sites) to assess if Sentinel-1 could be used for detecting selective logging activities near the legal limit within the Brazilian Legal Amazon. Unfortunately RADARSAT-2 and PALSAR-2 imagery did not cover the highest intensity logging sites, so we could not perform equivalent analyses with these datasets. RF classification and validation was performed on this subset of the Sentinel-1 data in the manner detailed above for the full dataset.

### 3.2. Time series analyses

We tested whether a time series of Sentinel-1 data displayed discernible changes in pixel values after selective logging with the BFAST algorithm (Verbesselt et al., 2012, 2010) in program R (R Core Team, 2018). BFAST estimates the timing of abrupt changes within a time series (breakpoint hereafter) and has been successfully utilized with a range of data types (e.g. Landsat, MODIS, SAR, etc.). The metrics used in searching for breakpoints in the full Sentinel-1 time series (approximately 55 scenes from October 2016 – August 2018) were the two most important predictor variables identified from RF models. The limited temporal coverage of RADARSAT-2 and PALSAR-2 at our study sites precluded time series analyses with these datasets. BFAST was used to assess if a suitable model with one or no breakpoints was appropriate and included tests for coefficient and residual-based changes in the expected value (i.e. the conditional mean). Where breakpoints were identified, we determined if they coincided with the timing of selective logging activities (June – October) and regarded these as true detections. Breakpoints in unlogged areas and breakpoints outside the timing of logging activities were considered false detections. In addition, the relationship between the frequency of breakpoints within an FMU and its logging intensity was examined to understand potential thresholds in logging intensity above which variables could be used to monitor selective logging activities through time series analyses.

Finally, we examined if the relationship between logging intensity and the rate of detections and false alarms was consistent between logging locations (i.e. a scattered subset of pixels in an area) and an entire region (i.e. all pixels within a bounding box). The timing of breakpoints was mapped for two 500 m X 500 m test regions within the Saraca study area (one logged and one unlogged). A limited number of small test regions were chosen because of the computationally expensive nature of the pull request in Earth Engine (e.g. two 1 km regions query > 1 million records for export). Only breakpoints during the time period associated with logging were mapped (June – October).

## 4. Results

### 4.1. Random Forest classification of selective logging

The single-image detection results for all sensors revealed that in order to get false discovery rate (FDR) values sufficiently low (e.g. 10-20%), the corresponding detection rates (DR) of selective logging were of almost no value (< 5%) for reliably forest monitoring. In general, the following results suggest that regions that have experienced selective logging do not show consistent differences from unlogged areas in the metrics we used for classification. The second analysis (section 4.2) therefore deals with detection of selective logging with time series data and provides better results.

#### 4.1.1. Sentinel-1

Random Forest detection performance for Sentinel-1 is shown in Figure 3 (top). Both the detection and false alarm rates were close to 1 until the threshold exceeds ∼0.4, meaning almost every pixel in an image would be detected as logged. This suggests difficulty distinguishing logged and unlogged observations, and many unlogged observations were being misclassified as logged (Figure S1). In general, the detection, false alarm, and false discovery rates (across the range of threshold values) were insufficient for reliable classification of selective logging with Sentinel-1 data at the intensities within our study areas (6-25 m^3^ ha^−1^). For example, even if a FDR of 30% were acceptable, this would yield a detection rate < 20%, which would be of little practical value. Thus, attempts to strongly limit the false discovery rate (commission error of logged observations) would require a high threshold value and result in very few detections. Overall, this suggests that using single images from Sentinel-1on their own to detect and map selective logging activities would be fraught with error with the classification approach used here.

**Figure 3.**
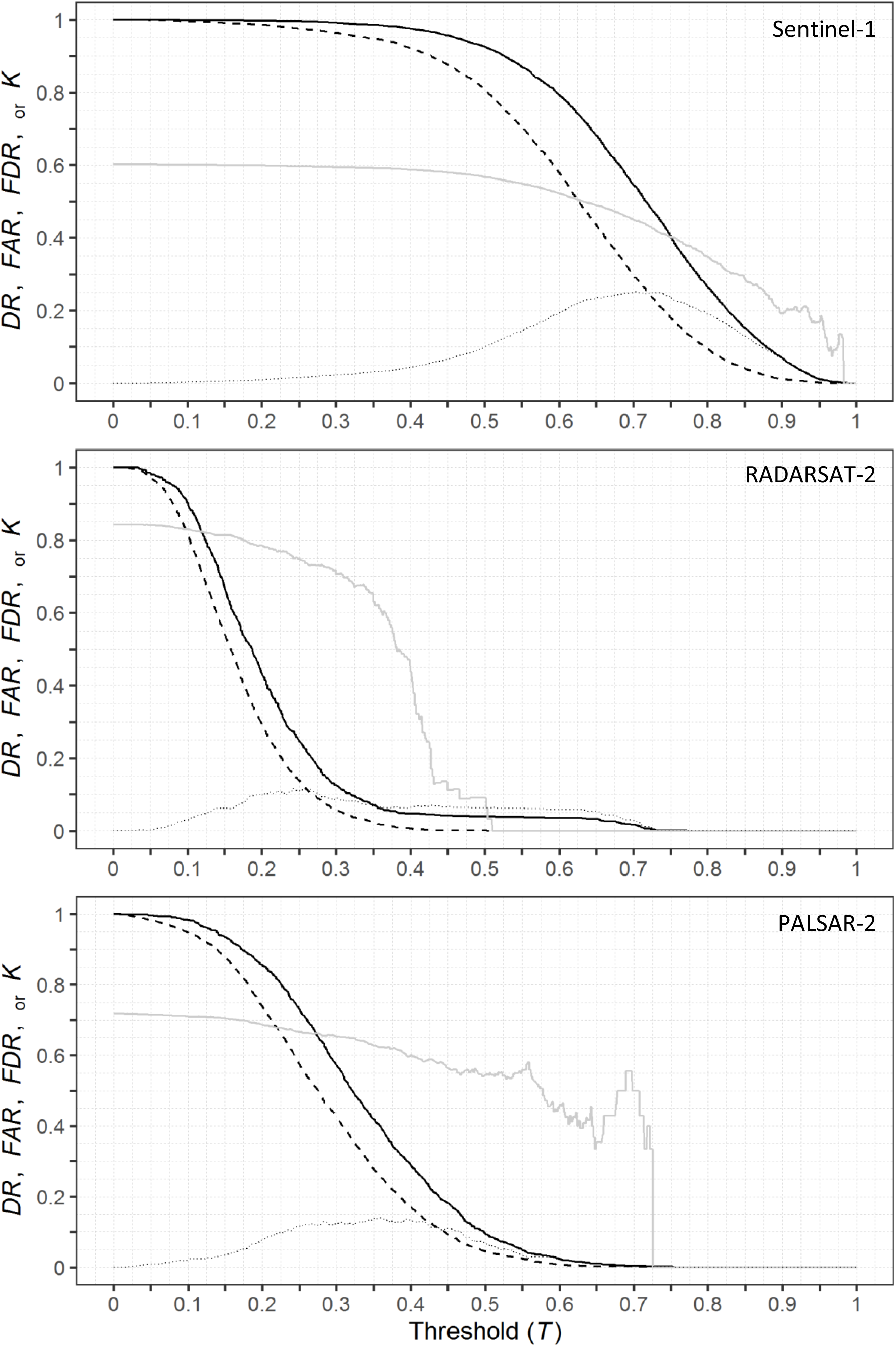
Random Forest model performance across the range of threshold values (*T*) for classification with SAR data. The Detection Rate (DR) and False Alarm Rate (FAR) are the solid and dashed black lines, respectively. Also shown are the corresponding values of the False Discovery Rate (FDR) and Cohen’s kappa (*k*) as solid and dotted grey lines, respectively.

#### 4.1.2. RADARSAT-2

Random Forest performance for RADARSAT-2 is shown in Figure 3 (middle). Both the false alarm rate and the detection rate rapidly declined as the threshold value was initially increased, again suggesting difficulty in distinguishing logged and unlogged observations. In contrast to Sentinel-1, RADARSAT-2 was less likely to label an observation as logged and very few observations had likelihood values above 0.5 (Figure S2). It should be noted that the logging records that coincided with RADASAT-2 data were from a single FMU that was relatively low intensity (10 m^3^ ha^−1^). Consequently, the performance displayed here may not be a full appraisal of RADARSAT-2 capabilities. Given how poorly the model performed, however, it is uncertain that a vast improvement would occur with better training datasets. Overall, our results suggest that RADARSAT-2 data cannot be used to effectively monitor low-intensity selective logging activities using pixel-based differences between logged and unlogged areas. However, additional tests with data at higher logging intensities should be pursued.

#### 4.1.3. PALSAR-2

Random Forest classification performance for PALSAR-2 is shown in Figure 3 (bottom). In general, the performance of PALSAR-2 was equally poor at distinguishing logged and unlogged observations as RADARSAT-2 and Sentinel-1 (Figure S3). The final rise in the false discovery rate in Figure 3, before it drops to zero, is the result of calculating proportions from very small sample sizes (e.g. 5 of 10 observations predicted logged were actually unlogged). Similar to RADARSAT-2, the selective logging data that coincided with PALSAR-2 imagery was from two relatively low-intensity FMUs (9 - 11 m^3^ ha^−1^). Again, however, more data at higher logging intensities seems unlikely to improve classification performance to the desired level. For example when the data from Sentinel-1 was restricted to just the low intensity sites used in the PALSAR-2 analyses, there was effectively no change in the rates of detection and false discovery compared to the results from all logging intensities with Sentinel-1 (Figures S4 and Table S7). Thus, the lack of higher intensity logging data probably had little impact on the results for PALSAR-2. In general, this suggests that the limitations in distinguishing logged and unlogged pixels are inherent in the data and metrics we used for classification (for all three data sets).

#### 4.1.4. Sentinel-1 classification of high intensity logging

Detection performance of Sentinel-1 data for the highest intensity FMUs is shown in Figure 4. Despite limiting the detection task to the most intensively logged FMUs (as well as unlogged observations), the detection rate and false discovery rate values were comparable to the results that used the full range of logging intensities. Instead, improvement in model performance was associated with better discrimination of unlogged observations (i.e. compare the commission and omission errors for the unlogged class between Tables 2 and 5). Essentially, the model was able to better identify unlogged forest, presumably because the more “confusing” observations (i.e. the low intensity FMUs) were absent and could not muddle the distinction between logged and unlogged observations (Figures S5). Overall, our results suggest Sentinel-1 data cannot be used in the classification of pixel-based differences to monitor selective logging activities with reasonable precision, even at the most intensively logged regions within the Amazon.

**Figure 4.**
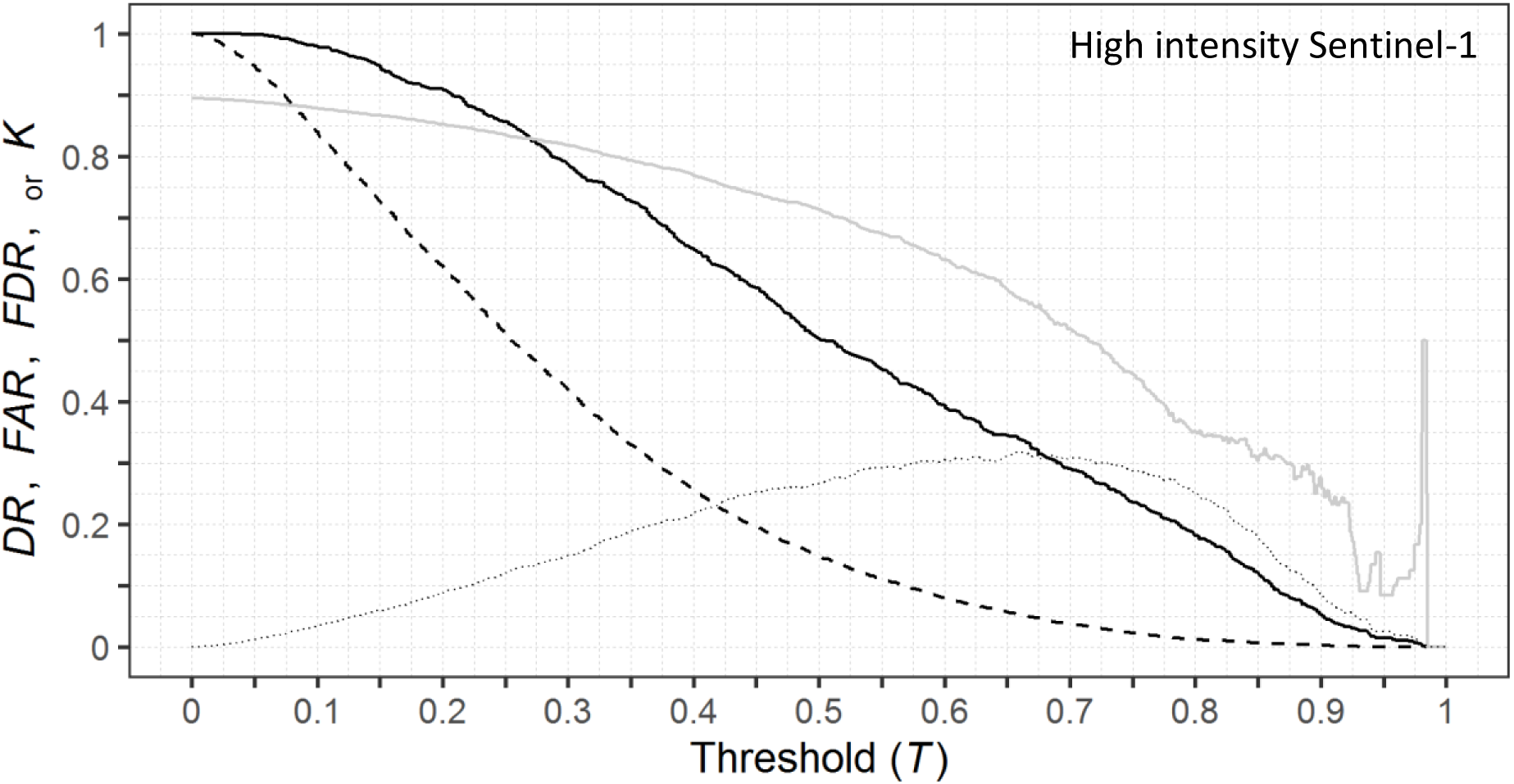
Random Forest model performance across the range of threshold values (*T*) for classification of Sentinel-1 data with a subset of the most intensively logged sites. The Detection Rate (DR) and False Alarm Rate (FAR) are the solid and dashed black lines, respectively. Also shown are the corresponding values of the False Discovery Rate (FDR) and Cohen’s kappa (solid and dashed grey lines, respectively).

### 4.2. Sentinel-1 time series analyses

The two most important predictor variables from the Sentinel-1 RF model were the Sum Average metric (Haralick 1973) on the VV and VH bands (Figure S6, Equation S1). A plot of VV sum average values through time for six randomly selected tree harvest locations at the Saraca site is shown in Figure 5 and suggests selective logging decreased the value of this metric. In addition, histograms of the timings associated with all breakpoints at three FMUs are shown in Figure 6 and indicates the time frame of the breakpoints mainly occurred within the logging season for those FMUs logged above 20 m^3^ ha^−1^. In contrast, the time periods associated with breakpoints at lower logging intensities were shifted toward the onset of the rainy season in late 2017 – early 2018, however, all FMUs showed an uptick in breakpoints associated with the rainy season (Figure 6). This suggests that Sentinel-1 time series data could be used to detect and monitor selective logging activities from areas that have experienced logging close to the legal limit in Brazil (30 m^3^ ha^−1^), particularly if the detection time-frame is narrowed to within the known logging season.

**Figure 5.**
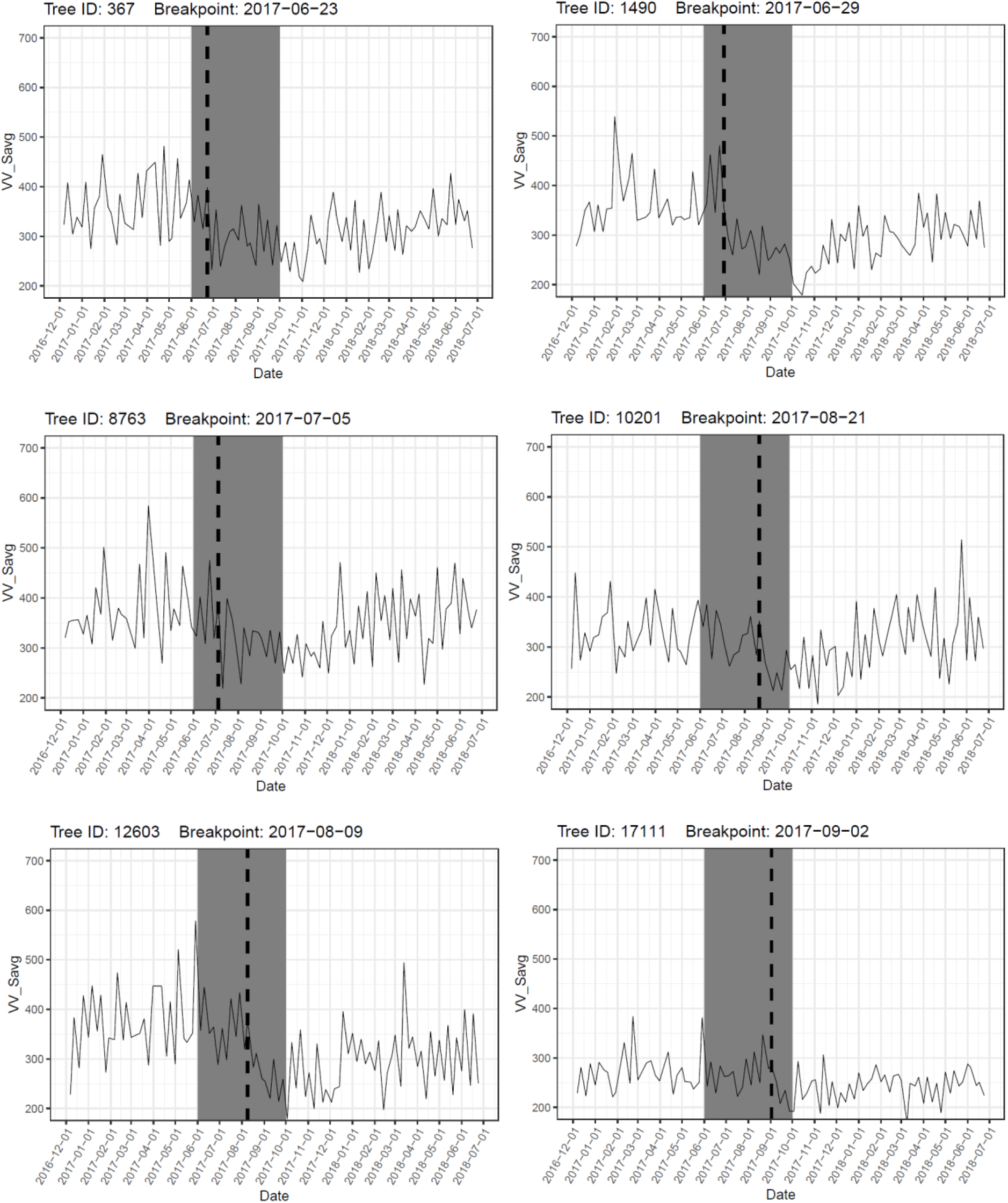
Breakpoint dates identified by the BFAST algorithm from six randomly selected points within the Saraca study region. The time series of the VV sum average texture measure is plotted in black, the selective logging period is shaded in grey, and the identified breakpoint date is labelled with a vertical dashed line.

**Figure 6.**
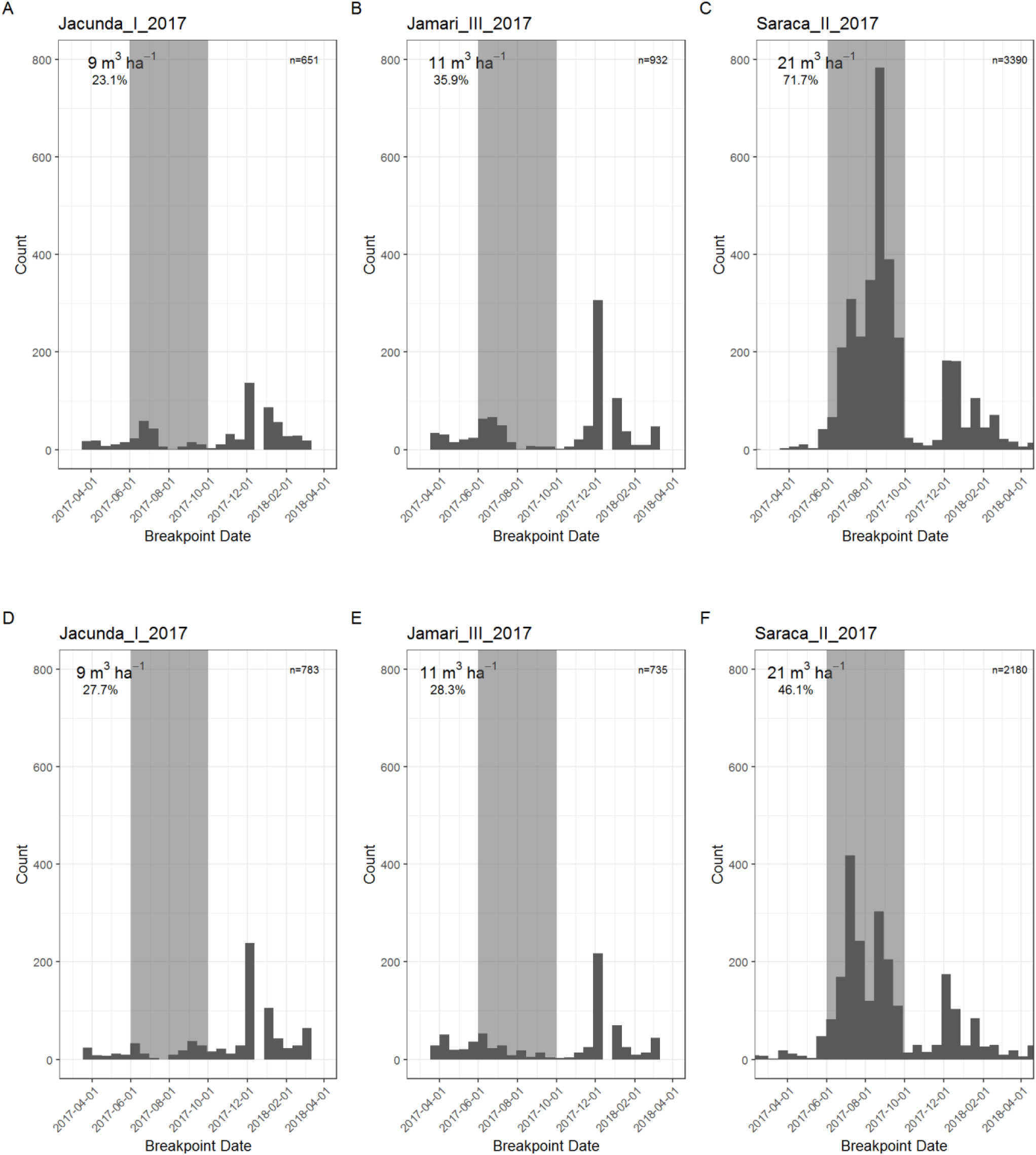
Histograms of breakpoint dates associated with time series analyses of the Sentinel-1 sum average texture measure for three study regions in the Brazilian Amazon for the VV (top row) and VH (bottom row) bands. The logging intensity and the proportion of observations with breakpoints in the data are in the upper left of each panel. The time period coinciding with logging activities is shaded in grey.

When the value of the VV sum average metric was monitored through time in pixels known to be logged and unlogged, the proportion of pixels with a significant breakpoint in their time series increased as the logging intensity of the FMU increased (Figure 7A). Approximately 70% of logged pixels in high logging intensity FMUs had a breakpoint, however, nearly 25% of unlogged pixels showed a breakpoint in their time series (i.e. 25% false alarm rate). This false alarm rate was generally consistent through logging intensities approaching 15 m^3^ ha^−1^ and suggests no signal in pixels logged at low to moderate intensities (Figure 7A). When the breakpoints were assessed only over the time period associated with logging (to remove the false peak associated with the rainy season), the relationship showed a similar pattern whereby the FMUs logged at the highest intensities showed a large rise in breakpoints above a background false alarm rate that was relatively constant up through moderate logging intensities (Figure 7B). At the highest intensities, the detection rate was > 50% and the false alarm rate was approximately 10%. These results further support the idea that FMUs logged at low to moderate intensities do not show a distinct time series signal whereas FMUs logged at higher intensities do. Overall, this suggests that FMUs logged at intensities closer to the legal limit within the Brazilian Legal Amazon (30 m^3^ ha^−1^) should show a noticeable spike in the number of breakpoints within its time series above a background false alarm rate and could be used to detect logging activities in the dry season.

**Figure 7.**
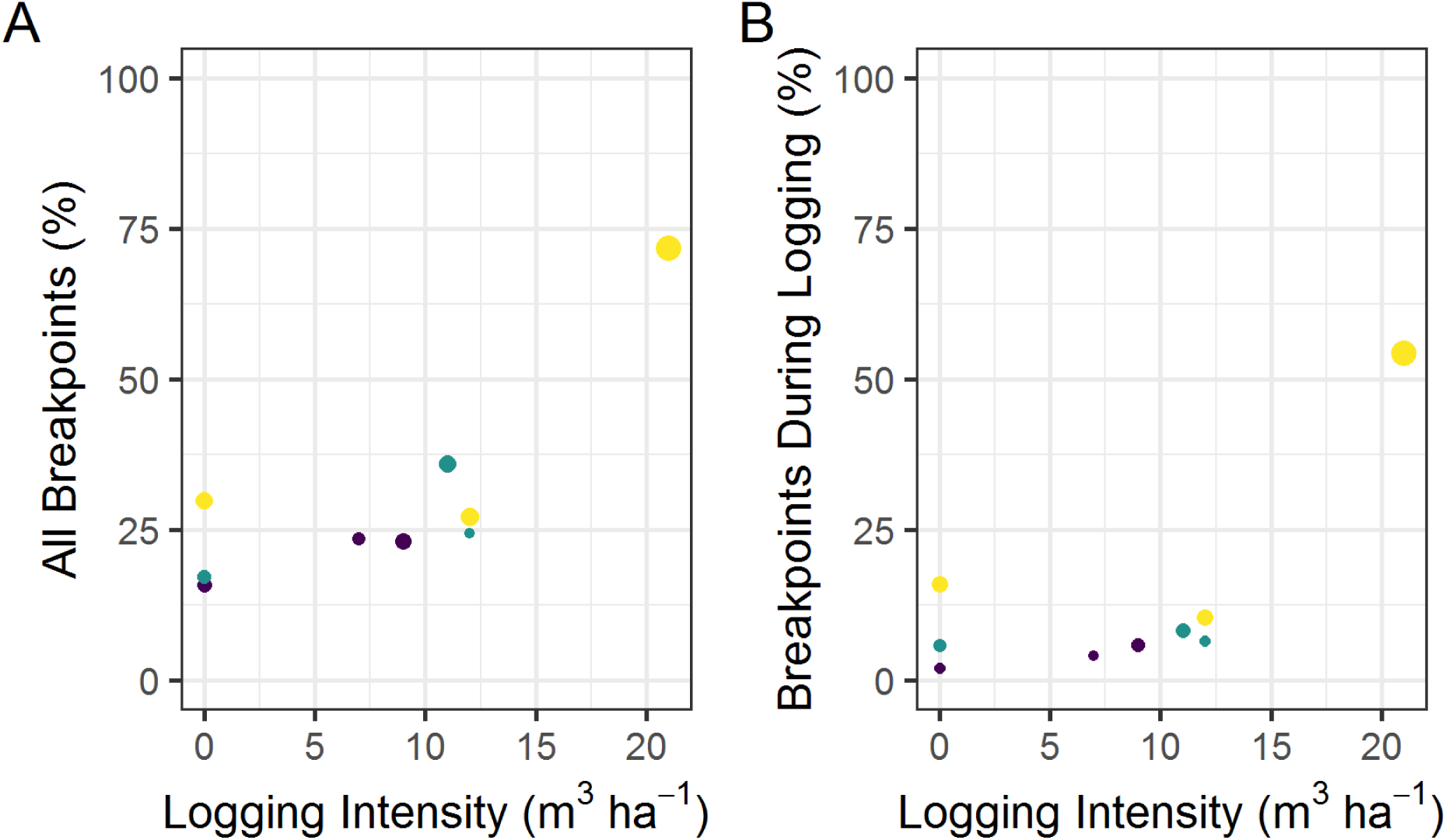
The relationship between the proportion of observation within a Forest Management Unit (FMU) that had a breakpoint identified within its Sentinel-1 VV sum average texture measure time series and the logging intensity of the FMU. The proportion of all observations (A) and the proportion that had a breakpoint that coincided with the logging season (B) are shown separately. The circle size corresponds to number of observations at each FMU and yellow, green, and purple colors represent the Saraca, Jamari, and Jacunda sites, respectively. See the supplementary material for the same analyses with the second and third best metric from Random Forest (Figure S7).

Approximately 55% and 20% of pixels in the logged and unlogged test regions had a breakpoint during the logging season (Figure 8A and B). These values are generally in agreement with our prior results from the subset of pixels where trees were removed (see Figure 7B). While 55% of the pixels in the logged test region did not have a tree removed, selective logging is associated with forest disturbances that go beyond the individually logged pixels (e.g. canopy gaps, skid trails, logging roads, etc.) and additional detections are expected. Only about 5% of the pixels in the logged test region were actually logged, however, it is clear from the Planet imagery (Figure 8C and D; Planet Team 2017) that more than 5% of the forest patch was disturbed by logging activities. Given the false alarm rate was around 20%, the difference between detections and false alarms might represent a value comparable with the amount of forest disturbance expected at this intensity (i.e. about 30%).

**Figure 8.**
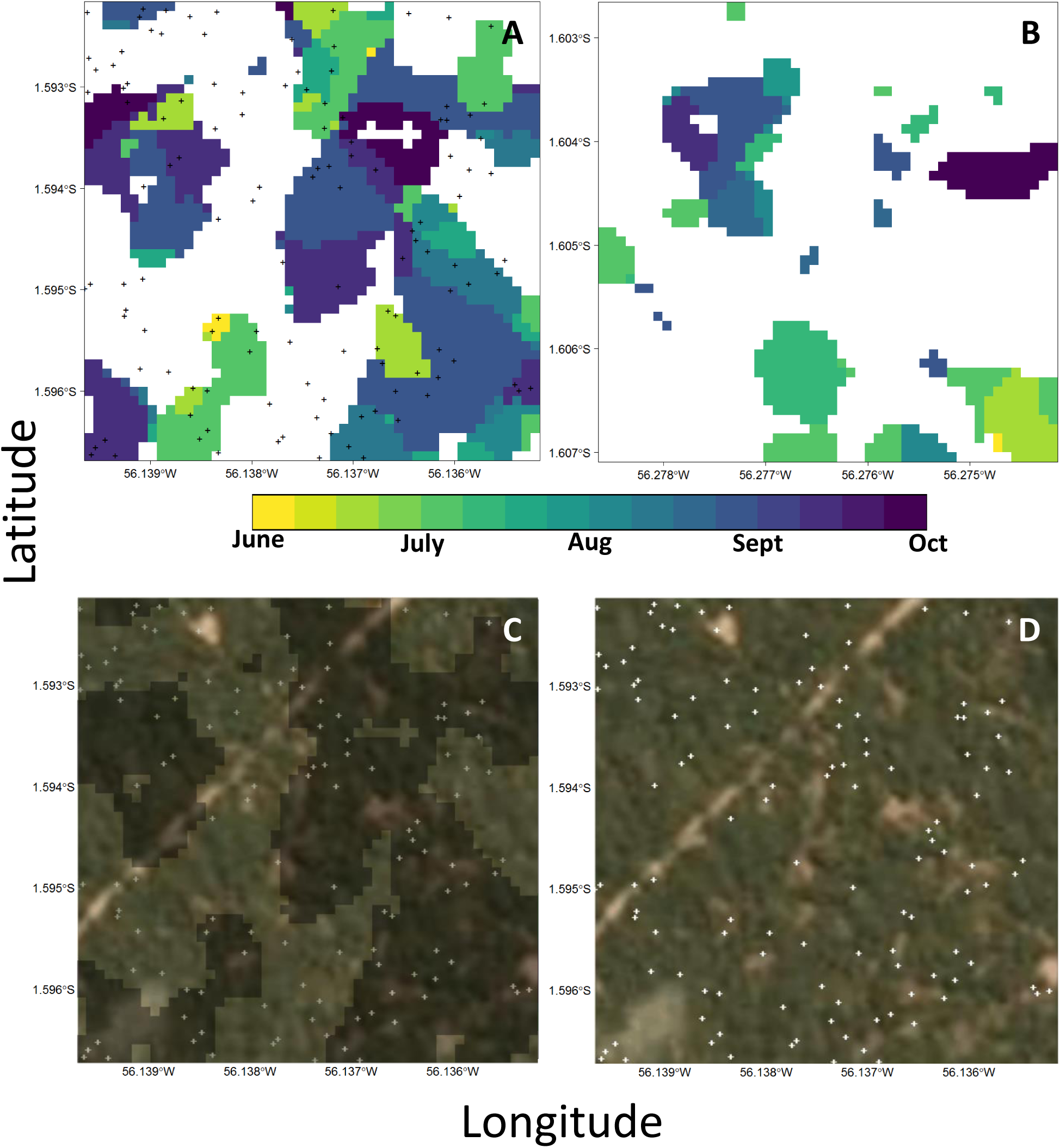
Map of predicted breakpoint dates for two 500m X 500m test regions, one logged (A) and one unlogged (B), in the Saraca National Forest, Para, Brazil. Logged tree locations are black crosses and the date of the breakpoint for each pixel is color coded by week, with white representing no breakpoint. Planet imagery (3 m) from 28 August 2017 overlaid with and without breakpoint locations (C and D) for the logged area (trees in white). Approximately 54% and 21% of the pixels in the logged and unlogged regions had breakpoints, respectively.

## 5. Discussion

We present the first multi-sensor comparison of SAR data for monitoring a range of selective logging intensities in the tropics. We demonstrated that L-band PALSAR-2, C-band RADARSAT-2, and C-band Sentinel-1 data performed inadequately at detecting tropical selective logging when using pixel-based attributes for classification. However, when analysing a time series of Seninel-1 texture measures, logged pixels displayed a strong tendency for a breakpoint in their time series as the logging intensity of the FMU increased. Moreover, the timing associated with the identified breakpoint generally coincided with active logging at the highest logging intensities. Overall, our results suggest that Sentinel-1 data could be used to monitor the most intensive selective logging, but a time series approach would be required to detect change. A number of studies have used Sentinel-1 time series data to monitor deforestation (Bouvet et al., 2018; Reiche et al., 2018a, 2018b), often in combination with optical data, however our study is the first to show it has the potential to be used exclusively to monitor selective logging.

### 5.1. Variable importance

In a number of cases the most important predictor variables from RF models involved the co-polarized channel (Figure S1), despite the generally accepted view that the cross polarized channel is best for detecting changes in forest cover (Joshi et al., 2016; Reiche et al., 2018a; Ryan et al., 2012; Shimada et al., 2014). The HH polarization of PALSAR-2 data has previously been shown to be sensitive to the early stages of deforestation, resulting from single-bounce scattering from felled trees (Watanabe et al., 2018). Our results support the idea that the co-polarized channel (for L- and C-band SAR) is useful and should not be ignored in forest disturbance detection analyses (e.g. Reiche et al., 2018a). While shorter wavelength SAR data, like C- and X-band, are known to be less sensitive to forest structure, because the radar signal mainly interacts with the forest canopy (Woodhouse, 2017; Flores-Anderson et al., 2019), the higher backscatter values in the co-polarized channel for all three sensors suggests predominantly rough surface backscattering from the forest canopy (as volume scattering generally results in roughly equal backscatter between co- and cross-polarized channels). This suggests that forest tracts subjected to more intensive selective logging than we studied (conventional logging permits with larger canopy gaps, large road networks, and many log landing areas) should possess a signal in the co-polarized channel that could be used to detect changes in canopy cover and should not be discarded (e.g. Reiche et al., 2018a).

Random Forest models offer an objective approach to selecting important variables for use in time series analyses. The Mean Decrease in Accuracy rankings were used to select the sum average texture measure in the time series results, corroborate their rankings (see Figures 7 and S7). The detection rate was highest with the best, lower with the second best, and lower still with the third. SAR data often has fewer bands than optical data, for example, so the choice of which metric to use in time series analyses may be more straightforward. However, many studies do not compare the results among metrics to select an optimal, relying instead on supposition (e.g. Reiche et al., 2018a). Our findings suggest Mean Decrease in Accuracy is useful for variable selection, even if the Random Forest models themselves are of little practical use (e.g. Figure 3).

### 5.2. Texture measures and detecting selective logging

In all cases the texture measures had the highest variable importance rankings (Figure S6). This corresponds with previous results with optical data, where detection of selective logging relied on the contextual information embodied within their calculation (Hethcoat et al. 2019). Similar to their results, the predictions of logging in our test areas were spatially correlated, presumably a consequence of the spatial window used in the calculation. Again, however, extra detections are expected from the accompanying forest disturbances associated with logging. Yet, in the context of accuracy assessment, an issue that has not received much attention within the remote sensing literature is how to report selective logging detections in the absence of robust field data on canopy gaps, roads networks, skid trails, log landing decks, etc. Others have shown that selective logging can be associated with 30-70% forest disturbance, despite the proportion of pixels having had a tree removed being closer to 10% (Asner et al., 2004, 2002; Putz et al., 2019), depending on the intensity and logging practices (reduced impact versus conventional). Clearly Figure 8A has false discoveries associated with the breakpoint detections, but some of the detections that do not occur at a tree location undoubtedly correspond with canopy gaps seen in the Planet imagery.

While the texture information clearly helped with detection of selective logging, a coherent understanding of what the sum average metric means, in terms of characterizing forest disturbances from selective logging or understanding the structural changes to forests associated with increasing and decreasing values, remains unknown. Attempts to generalize and interpret the meaning of textures have proven difficult over the years. However, some have suggested that high values in measures like variance, dissimilarity, entropy, and contrast were associated with visual edges whereas average, homogeneity, correlation, and angular second moment were associated with subtle irregular variations from continuous regions like forests or water (Hall-Beyer, 2017). More work is needed to understand the interpretation of textures measures that are so often employed in remote sensing classifications.

### 5.3. Combining sensors for classification

We chose not to combine any of the data types used here, partly because the inconsistent spatial and temporal coverage precluded such an analysis, but also because we wanted to assess the detection capabilities of each sensor on its own. Methods that combine data from multiple sensors (both other SAR platforms and/or optical data from Landsat or Sentinel-2) would likely perform better, corresponding with results for monitoring deforestation (Mercier et al., 2019; Reiche et al., 2018b, 2016, 2015). Indeed, prior work with Landsat data has shown strong detection of selective logging at similar intensities (Hethcoat et al., 2019), yet this work sought to establish a baseline with the SAR sensors available. The general direction and momentum for the advancement of detecting subtle forest disturbances from spaceborne SAR will likely require time series, polarimetric, and data fusion approaches, particularly in light of our findings that pixel-based differences between logged and unlogged areas with SAR backscatter alone cannot do the job effectively.

### 5.4. Longer time series in the tropics

Sentinel-1A began acquiring imagery regularly (approximately every 12 days) in late 2016 for most of Brazil, with Sentinel-1B following in late 2018. Consequently, a time series assessment was only possible for a single calendar year (roughly 2017) with the logging data sets we had access to. The BFAST algorithm is generally flexible and can be tuned with a baseline period if sufficient data are available, enabling assessments of longer and more variable time series (Verbesselt et al., 2010). The limited time series available is likely the reason many breakpoints for the less intensively logged sites occurred in December, presumably with the onset of the rainy season in earnest and an uptick in backscatter associated with moisture. Our analysis, however, was limited to a simpler test of one or no breakpoints – future work should explore how longer time series might improve detection of lower intensity logging, where seasonal patterns in backscatter can be established as a baseline to help reduce false alarms.

## 6. Conclusion

Tropical selective logging is fundamentally connected to global climate, biodiversity conservation, and human wellbeing (Lewis et al., 2015). Selective logging is often the first disturbance to affect primary forest (Asner et al., 2009), with road networks and ease of access facilitating further disturbances (e.g. increased fires, hunting or illegal logging). Efforts to detect and map selective logging with Sentinel-1, because of its global coverage and anticipated continuation missions (i.e. Sentinel-1C and D), are urgently needed to understand the capabilities this data stream might offer at advancing detection of tropical selective logging activities. With the successful launch of SAOCOM 1A in late 2018, the planned continuation of Sentinel-1 (with C and D), the opening of the ALOS PALSAR-1 archives, and the anticipated launches of SAOCOM 1B in 2019 and NISAR in 2021, an immense volume of freely available C- and L-band SAR data will, hopefully, usher in a new era of forest monitoring from space with SAR data. Our findings suggest that time series methods should be effective at detecting the most intensive selective logging in the Amazon with these data sets. Moreover, if a distinct dry season is characteristic of the study region, focusing detecting during this time frame can further bolster detection by removing false positive detections associated with seasonal rainfall.

## Acknowledgements

MGH was funded by the Grantham Centre for Sustainable Futures. JMBC was funded was funded by the Natural Environment Research Council (Agreement PR140015 between NERC and the National Centre for Earth Observation). RADARSAT-2 imagery was provided by MDA through an ESA agreement under proposal 126091 and PALSAR-2 imagery was provided by JAXA. We would like to thank Planet Labs for access to imagery through the education and outreach program.

