## Supplemental Material for "Detecting tropical selective logging with SAR data requires a time series approach"

**Table S1.** Acquisition dates for Sentinel-1, RDARSAT-2, and PALSAR-2 imagery used in classificication of selective logging.

| Sensor | Dates |
| --- | --- |
| Sentinel-1 | 2016-02-15 |
| | 2016-09-27 |
| | 2016-09-30 |
| | 2017-08-20 |
| | 2017-08-22 |
| | 2017-08-29 |
| | 2018-08-15 |
| | 2018-08-29 |
| RADARSAT-2 | 2012-08-19 |
| PALSAR-2 | 2016-09-06 |
| | 2017-07-05 |

**Table S2.** Cross-validation results for the number of of trees (*k*) and the number of variables to use at each node (*m*) that minimized the out-of-bag error rate on each training dataset.

| Sensor | nTree (*k*) | mTry (*m*) |
| --- | --- | --- |
| Sentinel-1 | 600 | 1 |
| RADARSAT-2 | 700 | 5 |
| PALSAR-2 | 700 | 4 |
| Sentinel-1 subset | 800 | 1 |

**Table S3.** Confusion matrix summarizing Random Forest (RF) model classifications of logged and unlogged observations at three study areas in the Brazilian Amazon, derived from Sentinel-1 data. Data were split into 75% training and 25% validation. Matrix numbers are pixel counts with the validation data (n = 13,401). The classification threshold (*T*) for RF models was set to maximize Cohen's kappa. The corresponding values for overall accuracy (OA), the false discovery rate (FDR), and the detection rate (DR) are provided against the validation dataset.

| **Sentinel -1**<br>OA: 64.3%<br>$\kappa$: 0.25<br>FDR: 44.6%<br>DR: 53.5% | | Reference Class<br>Logged | Reference Class<br>Unlogged | ***T* = 0.71**<br>Commission Error (%) |
|---|---|---|---|---|
| Predicted Class | Logged | 2861 | 2299 | 44.6 |
| | Unlogged | 2489 | 5752 | 30.2 |
| | Omission Error (%) | 46.5 | 28.6 | |

**Table S4.** Confusion matrix summarizing Random Forest (RF) model classifications of logged and unlogged observations at two study areas in the Brazilian Amazon, derived from RADARSAT-2 data. Data were split into 75% training and 25% validation. Matrix numbers are pixel counts with the validation data (n = 4,903). The classification threshold (*T*) for RF models was set to maximize Cohen's kappa. The corresponding values for overall accuracy (OA), the false discovery rate (FDR), and the detection rate (DR) are provided against the validation dataset.

| **RADARSAT-2**<br>OA: 75.6%<br>$\kappa$: 0.12<br>FDR: 75.0%<br>DR: 27.4% | | Reference Class<br>Logged | Reference Class<br>Unlogged | ***T* = 0.24**<br>Commission Error (%) |
|---|---|---|---|---|
| Predicted Class | Logged | 211 | 643 | 75.0 |
| | Unlogged | 559 | 3490 | 13.8 |
| | Omission Error (%) | 72.6 | 15.4 | |

**Table S5.** Confusion matrix summarizing Random Forest (RF) model classifications of logged and unlogged observations at two study areas in the Brazilian Amazon, derived from PALSAR-2 data. Data were split into 75% training and 25% validation. Matrix numbers are pixel counts with the validation data (n = 4,122). The classification threshold (*T*) for RF models was set to maximize Cohen's kappa. The corresponding values for overall accuracy (OA), the false discovery rate (FDR), and the detection rate (DR) are provided against the validation dataset.

| **PALSAR-2** | | | | **$T$ = 0.36** |
|---|---|---|---|---|
| OA: 64.2% | | | | |
| $\kappa$: 0.14 | | Reference Class | | |
| FDR: 62.4% | | Logged | Unlogged | Commission |
| DR: 40.5% | | | | Error (%) |
| Predicted Class | Logged | 471 | 782 | 62.4 |
| | Unlogged | 692 | 2177 | 24.1 |
| | Omission Error (%) | 59.5 | 26.4 | |

**Table S6.** Confusion matrix summarizing Random Forest (RF) model classifications of the most intensively logged and unlogged observations at three study areas in the Brazilian Amazon, derived from Sentinel-1 data. Data were split into 75% training and 25% validation. Matrix numbers are pixel counts with the validation data (n = 7,431). The classification threshold (*T*) for RF models was set to maximize Cohen's kappa. The corresponding values for overall accuracy (OA), the false discovery rate (FDR), and the detection rate (DR) are provided against the validation dataset.

| **Sentinel-1 High subset** | | | | **$T$ = 0.67** |
|---|---|---|---|---|
| OA: 88.5% | | | | |
| $\kappa$: 0.32 | | Reference Class | | |
| FDR: 55.9% | | Logged | Unlogged | Commission |
| DR: 33.2% | | | | Error (%) |
| Predicted Class | Logged | 261 | 331 | 55.9 |
| | Unlogged | 524 | 6315 | 7.7 |
| | Omission Error (%) | 66.8 | 5.0 | |

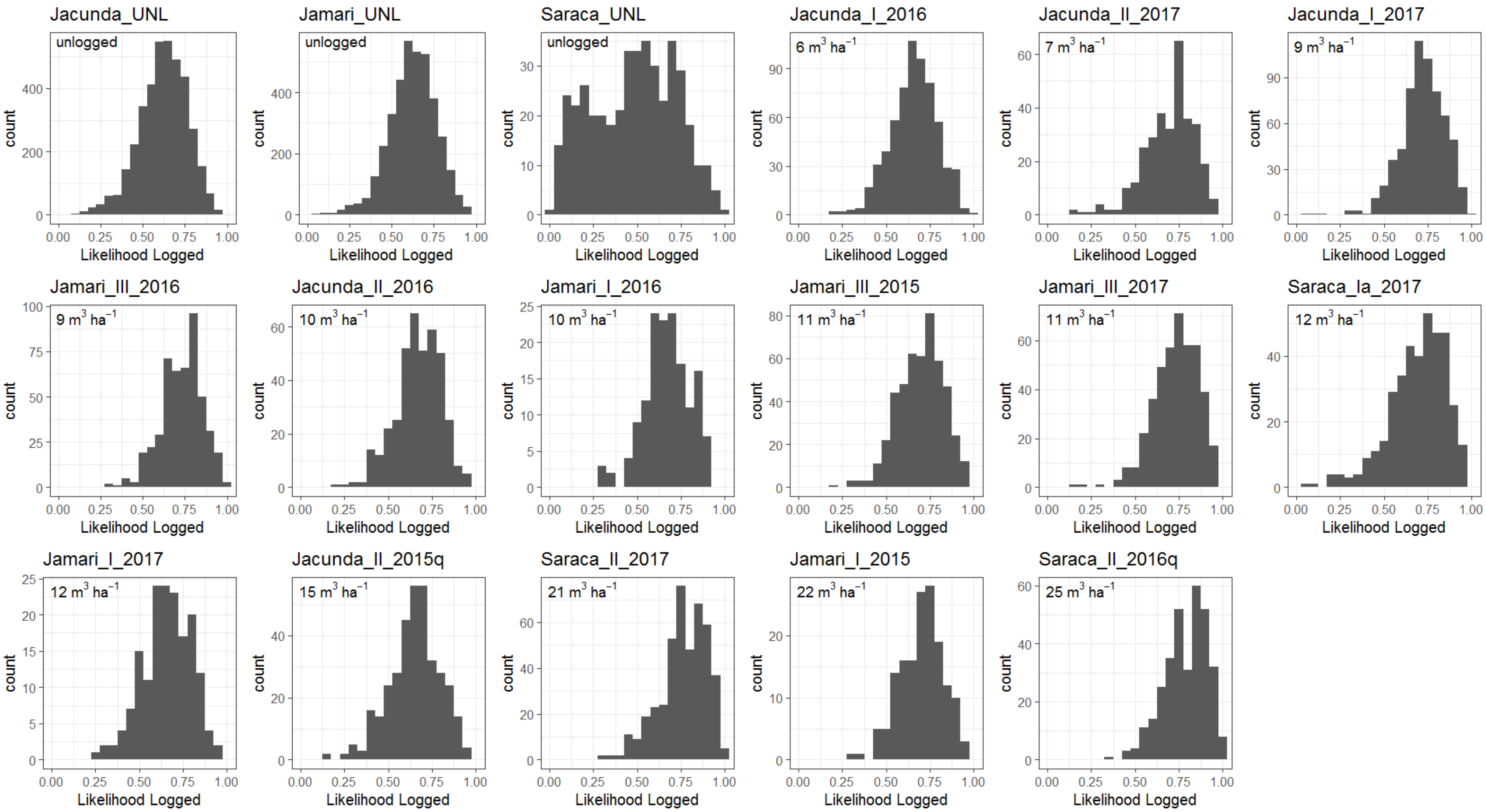


**Figure S1.** Histograms of the likelihoods (the proportion of votes for each class) for each observation with the full Sentinel-1 dataset (separated by FMU). The logging intensity is listed in the upper left of each panel.

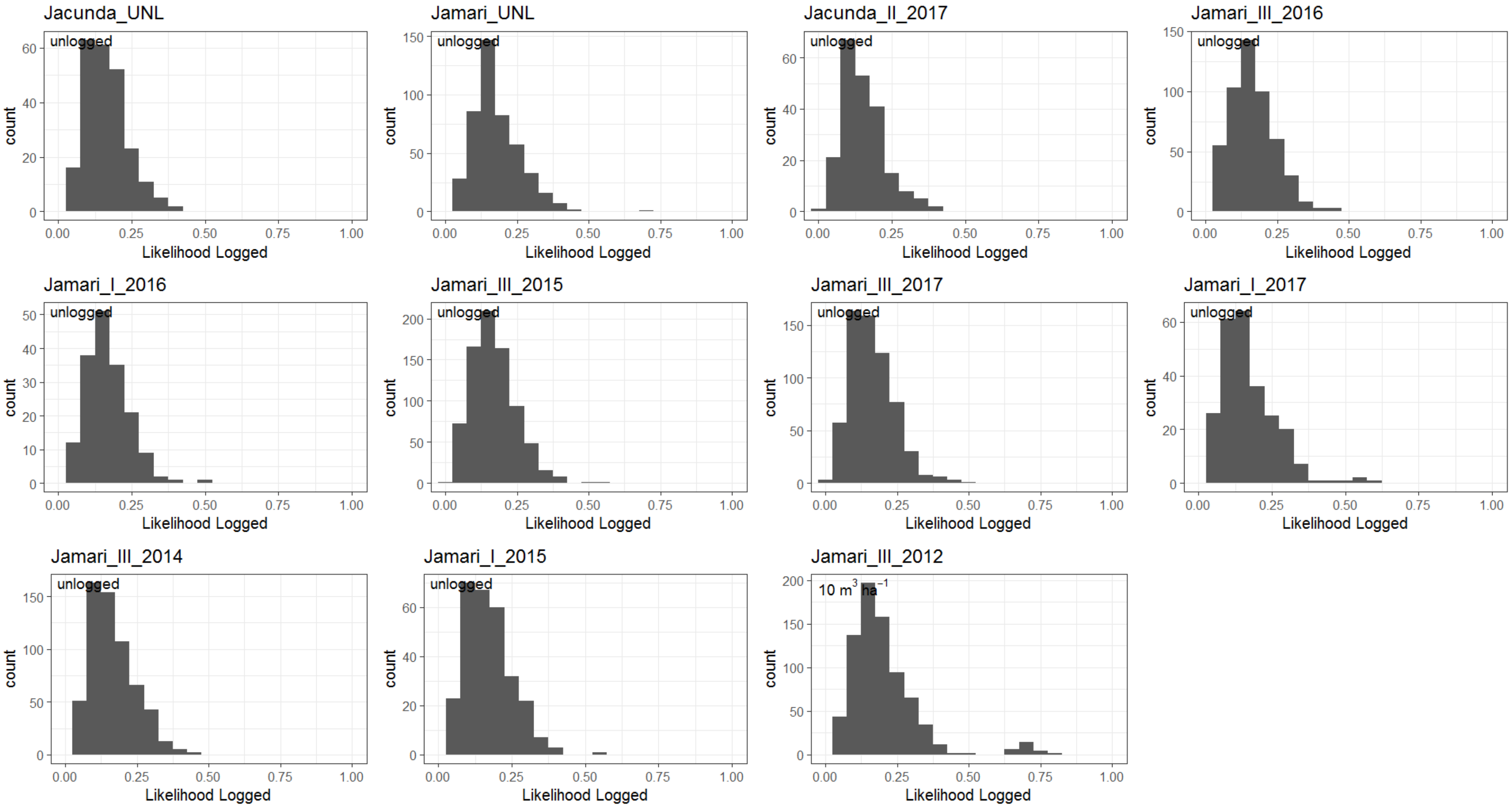


**Figure S2.** Histograms of the likelihoods (the proportion of votes for each class) for each observation with the RADARSAT-2 dataset (separated by FMU). The logging intensity is listed in the upper left of each panel.

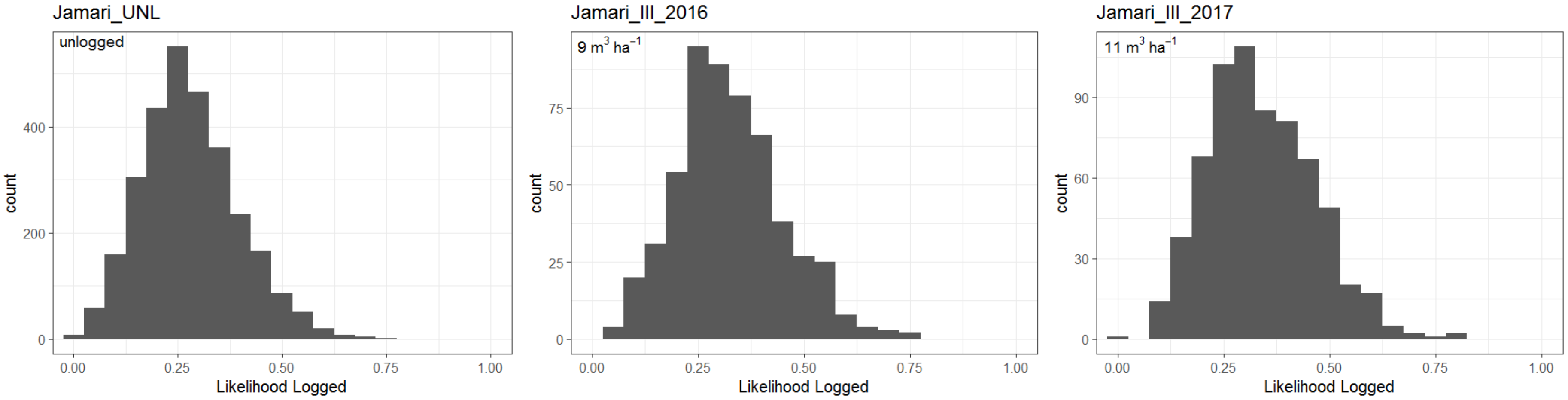


**Figure S3.** Histograms of the likelihoods (the proportion of votes for each class) for each observation with the PALSAR-2 dataset (separated by FMU). The logging intensity is listed in the upper left of each panel.

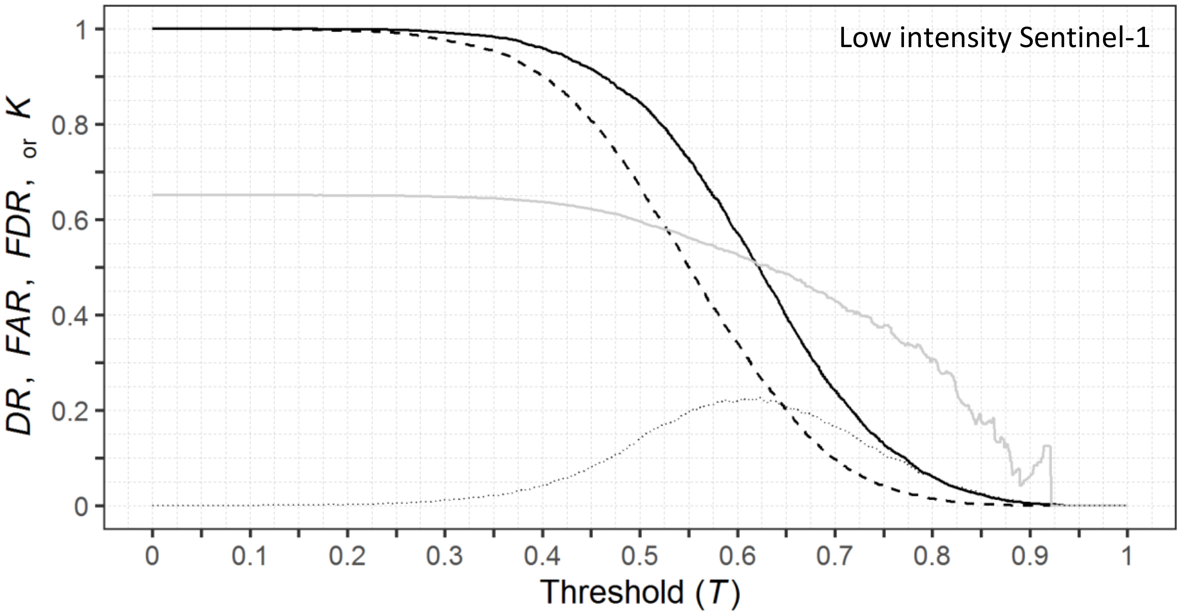


**Figure S4.** Random Forest model performance across all threshold values (*T*) for classification with Sentinel-1 (the same subset of low-intensity logging sites with that was used with the PALSAR-2). The Detection Rate (DR) and False Alarm Rate (FAR) are the solid and dashed black lines, respectively. Also shown are the corresponding values of the False Discovery Rate (FDR) and Cohen's kappa (*k*) as solid and dashed grey lines, respectively.

**Table S7.** Confusion matrix summarizing Random Forest (RF) model classifications of low-intensity logged and unlogged observations at two study areas in the Brazilian Amazon, derived from Sentinel-1 data (the same subset of sites used in the PALSAR-2 analyses). Data were split into 75% training and 25% validation. Matrix numbers are pixel counts with the validation data (n = 9,447). The classification threshold (*T*) for RF models was set to maximize Cohen's kappa. The corresponding values for overall accuracy (OA), the false discovery rate (FDR), and the detection rate (DR) are provided against the validation dataset.

| **Sentinel-1 Low subset** | | | | **$T$ = 0.62** |
|---|---|---|---|---|
| OA: 64.7% | | | | |
| $\kappa$: 0.23 | | Reference Class | | |
| FDR: 50.6% | | Logged | Unlogged | Commission |
| DR: 50.3% | | | | Error (%) |
| Predicted Class | Logged | 1659 | 1700 | 50.6 |
| | Unlogged | 1638 | 4450 | 26.9 |
| | Omission Error (%) | 49.7 | 27.6 | |

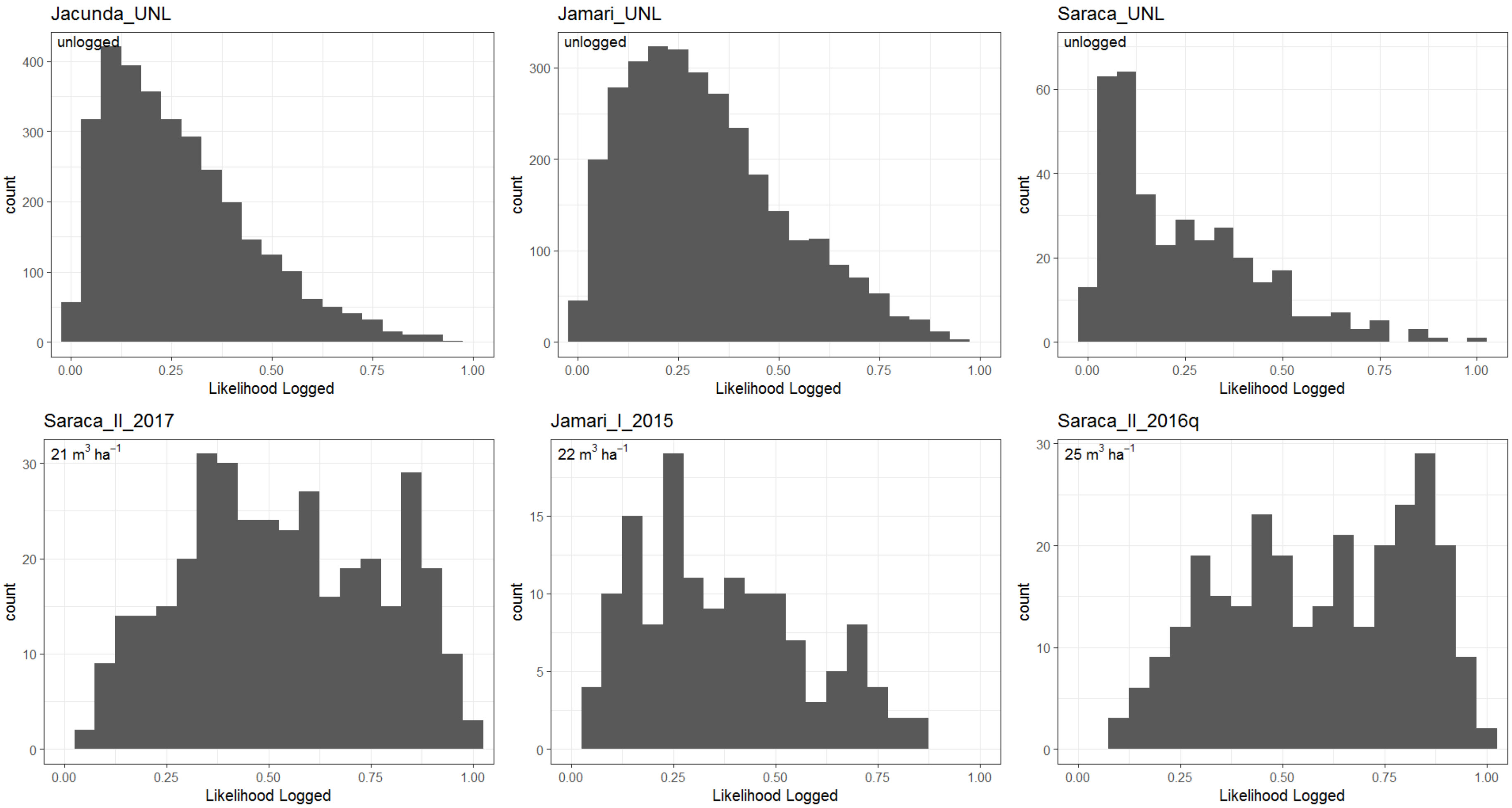


**Figure S5.** Histograms of the likelihoods (the proportion of votes for each class) for each observation with the subset Sentinel-1 dataset (separated by FMU). The logging intensity is listed in the upper left of each panel.

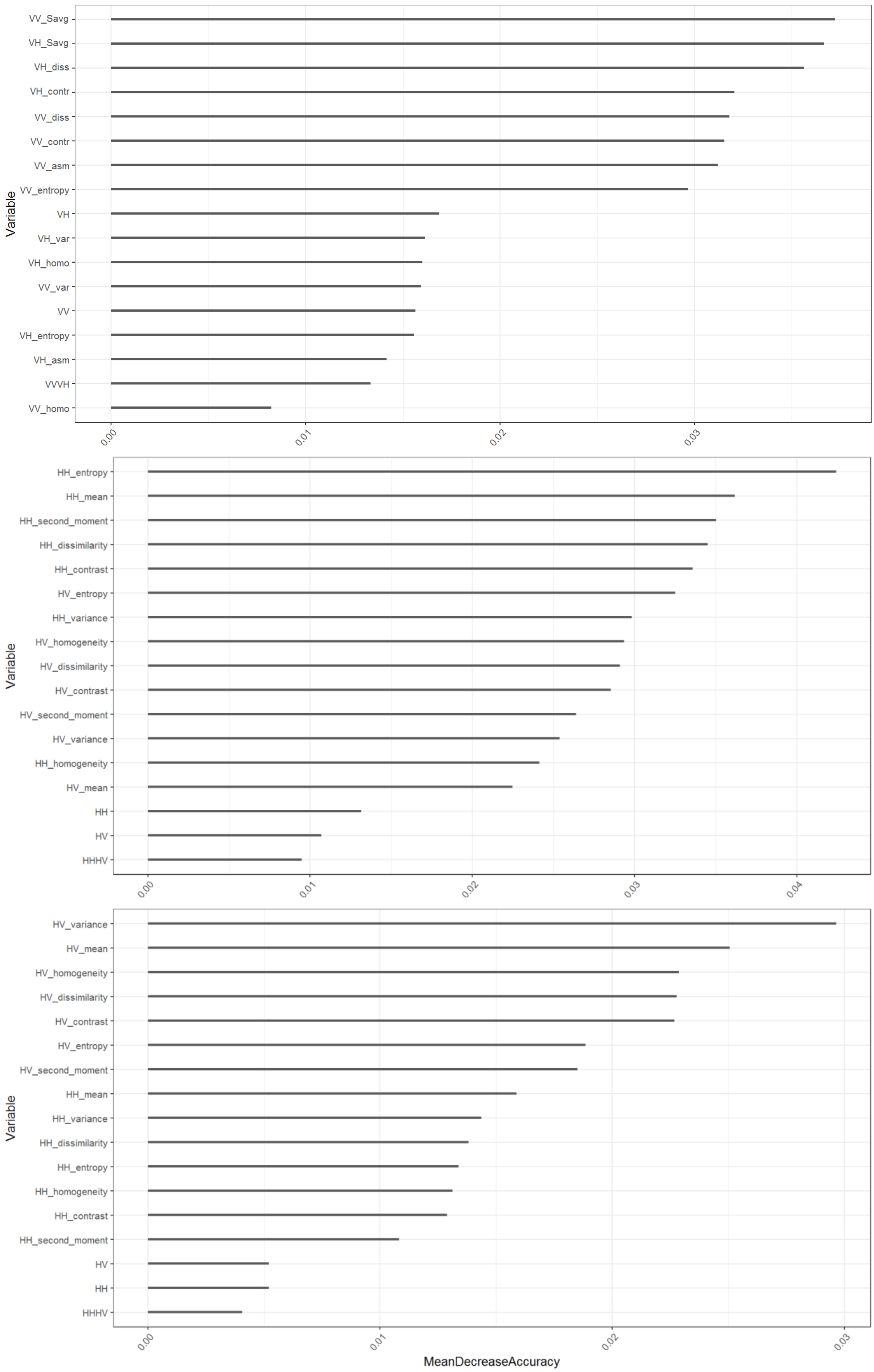


**Figure S6.** Random Forest model variable importance for Sentinel-1 (top), RADARSAT-2 (middle), and PALSAR-2 (bottom).

135

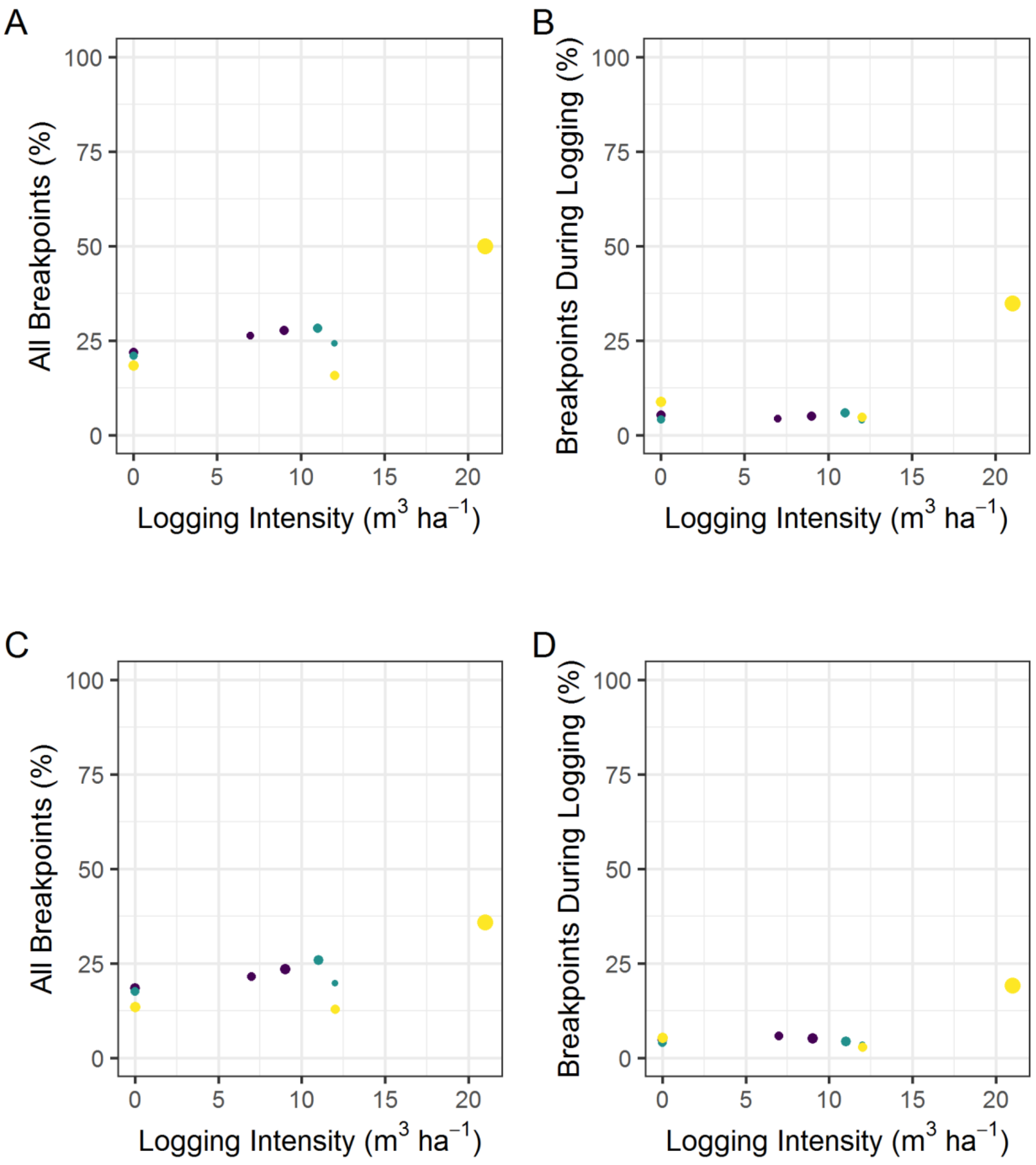


**Figure S7.** The relationship between the proportion of observation within a Forest Management Unit (FMU) that had a breakpoint identified within its Sentinel-1 sum average texture and dissimilarity measure time series and the logging intensity of the FMU for VH (bottom row). The proportion of all observations (A and C) and the proportion that had a breakpoint that coincided with the logging season (C and D) are shown separately. The circle size corresponds to number of observations at each FMU and yellow, green, and purple colors represent the Saraca, Jamari, and Jacunda sites, respectively.

Equation S1

$$Texture \sum Average = \sum_{i=2}^{2N_g} ip_{x+y}(i)$$

$N_g$: Number of distinct gray levels in quantized image

$x$ and $y$ are the coordinates (row and column) of an entry in the co-occurance matrix

$p_{x+y}(i)$ is the probability of co-occurances matrix coordinates summing to x+y
